# Pharmacokinetics Modeling and Simulation of Voriconazole Dosing and Safety in Patients with Liver Cirrhosis

**DOI:** 10.1101/563528

**Authors:** Taotao Wang, Miao Yan, Dan Tang, Yuzhu Dong, Li Zhu, Qian Du, Dan Sun, Jianfeng Xing, Yalin Dong

## Abstract

Voriconazole is used to treat invasive fungal disease and the optimal dose regimens are still unknown in cirrhotic Patients. The aim of this study was to determine the safety, to describe pharmacokinetics characteristics, and to optimize dosage regimens of voriconazole in cirrhotic patients. Data pertaining to voriconazole were collected retrospectively and analyzed using a population pharmacokinetics model. A total of 219 trough concentrations (*C*_min_) from 120 patients were analyzed. Voriconazole-related adverse events developed in 29 patients, with 69.0% of AEs developing within the first week after voriconazole treatment. The threshold *C*_min_ for AEs was 5.12 mg/L. A one-compartment model with first-order absorption and elimination adequately described the data. The Child-Pugh class was the only covariate in final model. Voriconazole clearance in patients with Child-pugh A and B cirrhosis (CP-A/CP-B) and Child-pugh C cirrhosis (CP-C) were 1.79 L/h and 0.99 L/h, respectively, the volume of distribution was 159.6 L, and the oral bioavailability was 91.8%. The elimination half-life was significantly extended for up to 61.8 – 111.7 h in cirrhotic patients. Model-based simulations showed that the appropriate maintenance doses are 75 mg/12 h and 150 mg/24 h intravenously or orally for CP-A/CP-B patients, and 50 mg/12 h and 100 mg/24 h intravenously or orally for CP-C patients. The results support voriconazole maintenance doses in LC patients should be reduced to one-fourth for CP-C patients and to one-third for CP-A/CP-B patients compared to that for patients with normal liver function. Monitoring *C*_min_ early could be a useful strategy to ensure the safety.

## 1. Introduction

Liver cirrhosis is a late stage of hepatic fibrosis and is a leading cause of death worldwide (1). Due to the immunocompromised state of cirrhotic patients and their requirement for use of steroids and broad-spectrum antimicrobial agents, as well as invasive procedures, invasive fungal disease (IFD) constitutes an important cause of morbidity in patients with liver cirrhosis (2-4). Voriconazole is a triazole antifungal agent with activity against a broad spectrum of clinically significant fungal pathogens, and it is used to treat and prevent IFD in clinical practice (5, 6).

Voriconazole is metabolized primarily by the *CYP2C19* isoenzyme while *CYP2C9* and *CYP3A4* contribute to a minor extent, and the large inter-and intra-individual variabilities of voriconazole pharmacokinetics are associated with *CYP2C19* genetic polymorphisms, liver dysfunction, drug-drug interaction and food intake (7-9). The voriconazole trough concentration (*C*_min_) is associated with treatment efficacy and safety (10, 11), and voriconazole *C*_min_ ranging from 1 – 5 mg/L was identified as the therapeutic target with appropriate clinical efficacy and safety (12). However, the reduced hepatic blood flow and enzyme activity in patients with liver cirrhosis can theoretically lead to voriconazole accumulation and make patients at risk of adverse events (AEs). Therefore, the voriconazole package insert recommends that while the standard loading dose regimens should be used, the maintenance dose should be halved in patients with Child-pugh A and B cirrhosis (CP-A/CP-B) (13). However, there is no recommended voriconazole regimen for patients with Child-pugh C cirrhosis (CP-C). Meanwhile, our previous study demonstrated that the halved dose still leads to a supratherapeutic level in the majority of cirrhotic patients, with a voriconazole *C*_min_ of 4.02 ± 2.00 mg/L (14, 15).

The present study aimed (1) to describe the pharmacokinetics characteristics and safety of voriconazole, and identify the factors influencing pharmacokinetics variability; and (2) to establish the optimal intravenous and oral voriconazole doses with an appropriate efficacy and safety profile in patients with liver cirrhosis.

## 2. Results

### 2.1 Patient characteristics

The demographic and clinical characteristics of the study population are presented in Table 1. The distributions of the initial voriconazole *C*_min_, as sampled before dosage adjustment, in relation to the different dosage regimens are shown in Figure 1. There were 40 CP-A/CP-B patients and 80 CP-C patients. Voriconazole *C*_min_ was measured in 219 samples (median of 1 per patient; range 1 – 7), and the median sampling time was 6 days (range 3 – 37 days) after the first dose. The initial voriconazole *C*_min_ was measured at a median time of 5 days (range, 3 – 27 days) after voriconazole was administered.

**Table 1.** Demographic and baseline clinical characteristics of patients

| Parameters | Group A |
| --- | --- |
| Child-pugh class (A:B:C) | 6:34:80 |
| Child-pugh score | 10.0 ± 1.8 (5 - 13) |
| MELD score | 18.3 ± 7.3 (6.2 – 40.1) |
| Demographic |  |
| Age (year) | 50.5 ± 13.2 (22 - 82) |
| Gender (male/female) | 95:25 |
| Weight (kg) | 63.1 ± 9.6 (40.5 - 88) |
| Type of cirrhosis, no. (%) of patients |  |
| Viral hepatitis | 93 (77.5%) |
| Alcoholic | 5 (4.2%) |
| Autoimmune hepatitis | 6 (5%) |
| Cholestasis | 2 (1.7%) |
| Hepatotoxic drugs | 5 (4.2%) |
| Cryptogenic | 9 (7.5%) |
| The laboratory data |  |
| RBC (10<sup>12</sup>/L) | 3.1 ± 0.7 (1.7 - 5.3) |
| WBC (10<sup>9</sup>/L) | 7.5 ± 4.8 (1.2 - 24.6) |
| PLT (10<sup>9</sup>/L) | 96.4 ± 85.6 (3 - 542) |
| ALT (U/L) | 101.3 ± 305.0 (3.0 - 3157.0) |
| AST (U/L) | 137.5 ± 277.0 (11.0 - 2718.0) |
| ALP (U/L) | 144.5 ± 100.3 (8 - 564) |
| Total bilirubin (μmol/L) | 243.7 ± 202.9 (5.1 - 901.6) |
| Albumin (g/L) | 31.9 ± 5.2 (19.1 - 45.4) |
| Serum creatinine (μmol/L) | 85.7 ± 77.5 (2.9 - 528.0) |
| INR | 1.9 ± 0.7 (1.0 - 4.2) |
| Concomitant medication, no. (%) of patients |  |
| *CYP2C19* inhibitor | 4 (3.3%) |
| *CYP2C19* inducer | 73 (60.8%) |
| *CYP3A4* inducer | 27 (22.5%) |

Note: continuous variables were expressed as mean ± standard deviation (range); ALT, alanine aminotransferase; AST, aspartate transaminase; ALP, alkaline phosphatase; INR, international normalized ratio.

**Figure 1.**
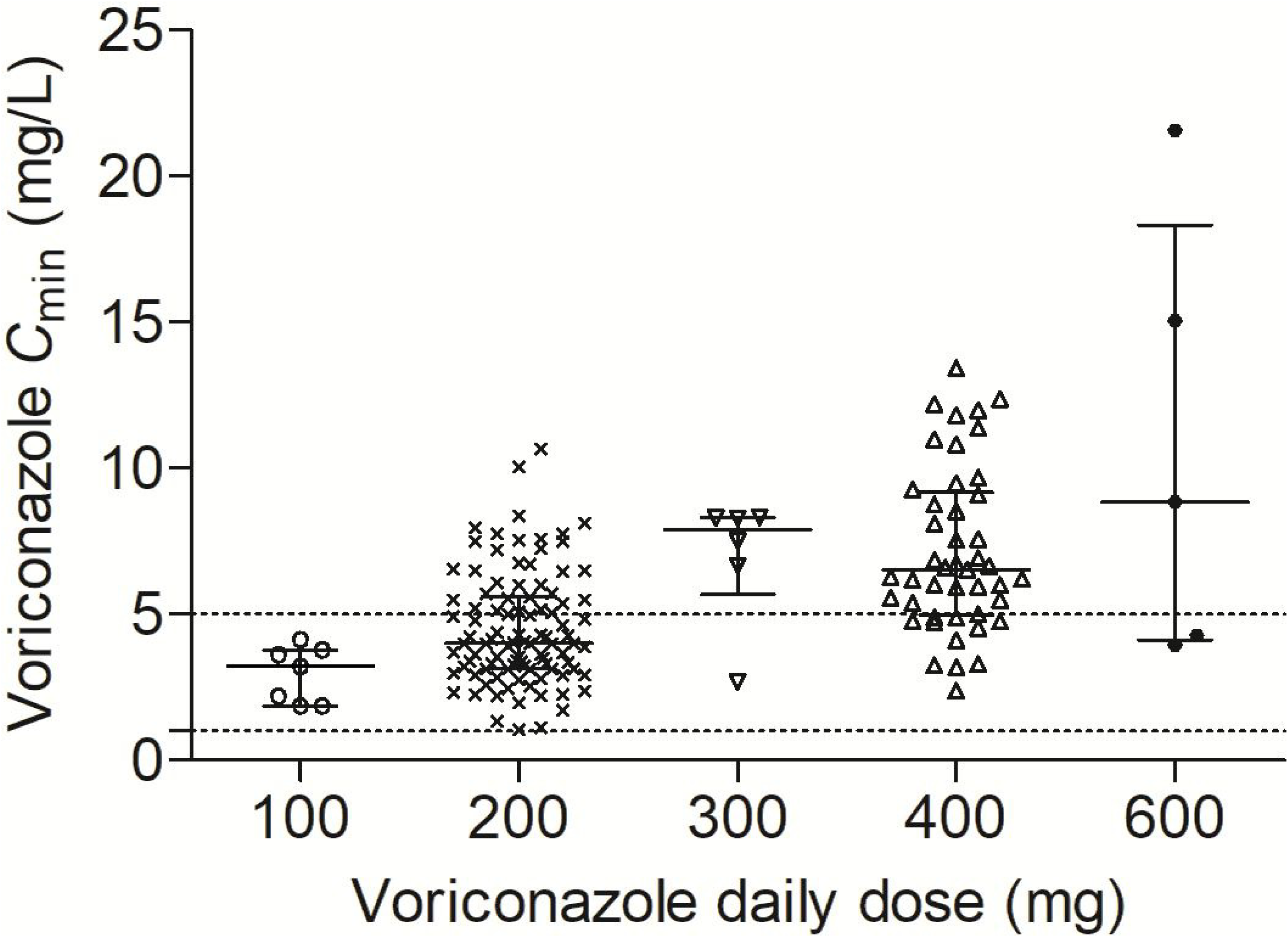
The distributions of the initial voriconazole *C*_min_ in relation to the different dosage regimens. The solid lines represent interquartile range (Q_25-75_). The dashed lines represent the voriconazole target trough plasma concentrations of 1 mg/L and 5 mg/L.

### 2.2 Voriconazole *C*_min_ and adverse events

Voriconazole-related AEs appeared in 29 (24.2%) patients. The most common AEs were neurotoxicity, visual disturbances and hallucination. The median voriconazole *C*_min_ in patients with AEs was 5.98 mg/L (range, 2.37 – 15.10 mg/L), and the median time from voriconazole treatment initiation to the development of AEs was 6 days (range, 2 – 15 days) (Table 2). Additionally, 69.0% (20/29) of voriconazole-related AEs developed within the first week after voriconazole treatment. The voriconazole *C*_min_ at the time of AEs emerging differed significantly from the initial *C*_min_ among patients without AEs (median 5.98 mg/L *vs* 4.76 mg/L, *P* = 0.045). The CART analysis showed that voriconazole-related AEs were more likely to occur in patients with *C*_min_ > 5.12 mg/L (*P* = 0.014). The incidence rates of AEs for patients achieving *C*_min_ values of >5.12 mg/L and ≤5.12 mg/L were 32.8% and 16.1%, respectively.

**Table 2.** Voriconazole trough concentrations and the time from voriconazole treatment initiation to the development of adverse events. Data were expressed as median (range).

| | Voriconazole $C_{\min}$ (mg/L) | Time (days) |
| --- | --- | --- |
| Visual disturbances (n = 6) | 4.69 (3.14 - 8.76) | 6 (2 - 8) |
| Hallucination (n = 3) | 3.25 (2.37 - 8.30) | 4 (2 - 4) |
| Neurotoxicity (n = 12) | 5.85 (3.30 - 15.10) | 7 (2 - 12) |
| Rash (n = 3) | 6.00 (5.14 - 10.65) | 4 (4 - 7) |
| Nausea and vomit (n = 2) | 4.12, 10.82 | 13, 15 |
| Hypokalemia (n = 2) | 6.71, 8.29 | 4, 8 |
| Nephrotoxicity (n = 1) | 6.17 | 7 |
| All patients | 5.98 (2.37 - 15.10) | 6 (2 - 15) |

$C_{\min}$ : trough concentration.

### 2.3 Population pharmacokinetics analysis

When each covariate was tested separately, the Child-Pugh class, toal bilirubin (TBIL), Child-Pugh score, international normalized ratio (INR), MELD score, gamma-glutamyl transferase (GGT), aspartate transaminase (AST), alkaline phosphatase (ALP), platelet (PLT), age, and alanine transaminase (ALT) showed significant impacts on the CL of voriconazole. The type of cirrhosis, body weight and CYP inducer or inhibitor did not significantly influence voriconazole parameters in this population. The incorporation of the Child-Pugh class as a covariate in the basic model on CL resulted in a significant reduction in OFV (△OFV = 28.1). The Child-Pugh class was the only significant covariate in the final PPK model. The parameters of final PPK model and bootstrap results are summarized in Table 3, and most of the parameters in the final model were reliably estimated and with relatively small RSEs. The model was associated with the variability of CL decreasing from 67.2% to 59.2%. The typical value and variability of CL in CP-A/CP-B and CP-C patients were calculated using an IF-THEN coding structure in the control model. The estimated typical values of CL for CP-A/CP-B and CP-C patients were 1.79 L/h and 0.99 L/h, respectively, and the corresponding average elimination *t*_1/2_ values were 61.8 h and 111.7 h.

**Table 3.** The pharmacokinetics parameters of basic model and final model and bootstrap results.

| Parameters | Basic model <sup>a</sup> |  | Final model <sup>b</sup> |  | Final model <sup>c</sup> |  | Bootstrap results |  |  |
| --- | --- | --- | --- | --- | --- | --- | --- | --- | --- |
|  | Typical | RSE | Typical | RSE | Typical value | RSE | Mean | 95%CI- | 95%CI- |
|  | value | (%) | value | (%) |  | (%) |  | Lower | Upper |
| CL (L/h) | 1.35 | 8.9 | 1.77 | 8.1 | CL<sub>1</sub> = 0.99 | 10.6 | CL<sub>1</sub> = 1.01 | 0.78 | 1.24 |
|  |  |  |  |  | CL<sub>2</sub> = 1.79 | 8.3 | CL<sub>2</sub> = 1.80 | 1.49 | 2.12 |
| V<sub>d</sub> (L) | 150.4 | 8.3 | 159.3 | 7.6 | 159.6 | 7.7 | 158.3 | 131.7 | 185.0 |
| k<sub>a</sub> (h<sup>-1</sup>) | 0.25 | 0.3 | 0.25 | 0.2 | 0.25 | 0.2 | 0.21 | 0.07 | 0.35 |
| F (%) | 91.2 | 7.3 | 91.8 | 6.6 | 91.8 | 6.7 | 92.4 | 79.7 | 105.1 |
| ω<sub>CL</sub> | 67.2% | 20.1 | 59.2 | 25.6 | ω<sub>CL1</sub> = 58.4 | 48.9 | ω<sub>CL1</sub> = 55.9 | 33.6 | 78.4 |
|  |  |  |  |  | ω<sub>CL2</sub> = 60.0 | 33.9 | ω<sub>CL2</sub> = 58.8 | 47.4 | 70.18 |
| ω<sub>vd</sub> | 63.3% | 32.8 | 65.8 | 27.1 | 66.4% | 27.1 | 65.2 | 54.2 | 76.3 |
| θ<sub>CPG</sub> | / | / | 0.43 | 14.1 | / | / | / | / | / |
| Proportional error (%) | 0.096 | 72.8 | 0.105 | 63.4 | 0.108 | 62.9 | 0.11 | 0.04 | 0.18 |
| Additive error (mg/L) | 1.10 | 25.8 | 1.03 | 29.6 | 1.03 | 30.4 | 0.98 | 0.59 | 1.37 |

CL, clearance; V<sub>d</sub>, the volume of distribution; k<sub>a</sub>, absorption rate constant; F, bioavailability;
<sup>a</sup> CL = θ<sub>1</sub> × e<sup>η<sub>1</sub></sup>; V<sub>d</sub> = θ<sub>2</sub> × e<sup>η<sub>2</sub></sup>; k<sub>a</sub> = θ<sub>3</sub>; F = θ<sub>4</sub>;
<sup>b</sup> CL = θ<sub>1</sub> × (1 - θ<sub>5</sub> × CPG)) × e<sup>η<sub>1</sub></sup>; V<sub>d</sub> = θ<sub>2</sub> × e<sup>η<sub>2</sub></sup>; k<sub>a</sub> = θ<sub>3</sub>; F = θ<sub>4</sub>;
<sup>c</sup> CL<sub>1</sub> = θ<sub>1</sub> × e<sup>η<sub>1</sub></sup> (for CP- C patients); CL<sub>2</sub> = θ<sub>2</sub> × e<sup>η<sub>2</sub></sup> (for CP- A/CP-B patients); V<sub>d</sub> = θ<sub>3</sub> × e<sup>η<sub>3</sub></sup>; k<sub>a</sub> = θ<sub>4</sub>; F = θ<sub>5</sub>.

The goodness-of-fit plots of models are shown in Figure 2. The final model described the data very well, where observed concentrations were close to the population and individual model-based predictions. The bootstrap analysis showed that for the final model, 786 out of 1000 runs converged successfully. The mean values and 95% confidence interval (CI) of parameters from bootstrap results are presented in Table 3. The parameters of the final PPK model were close to the mean bootstrap values and all included in the 95% CI. The VPC showed that most of the observed data were fell within the 90% prediction interval (Figure S1).

**Figure 2.**
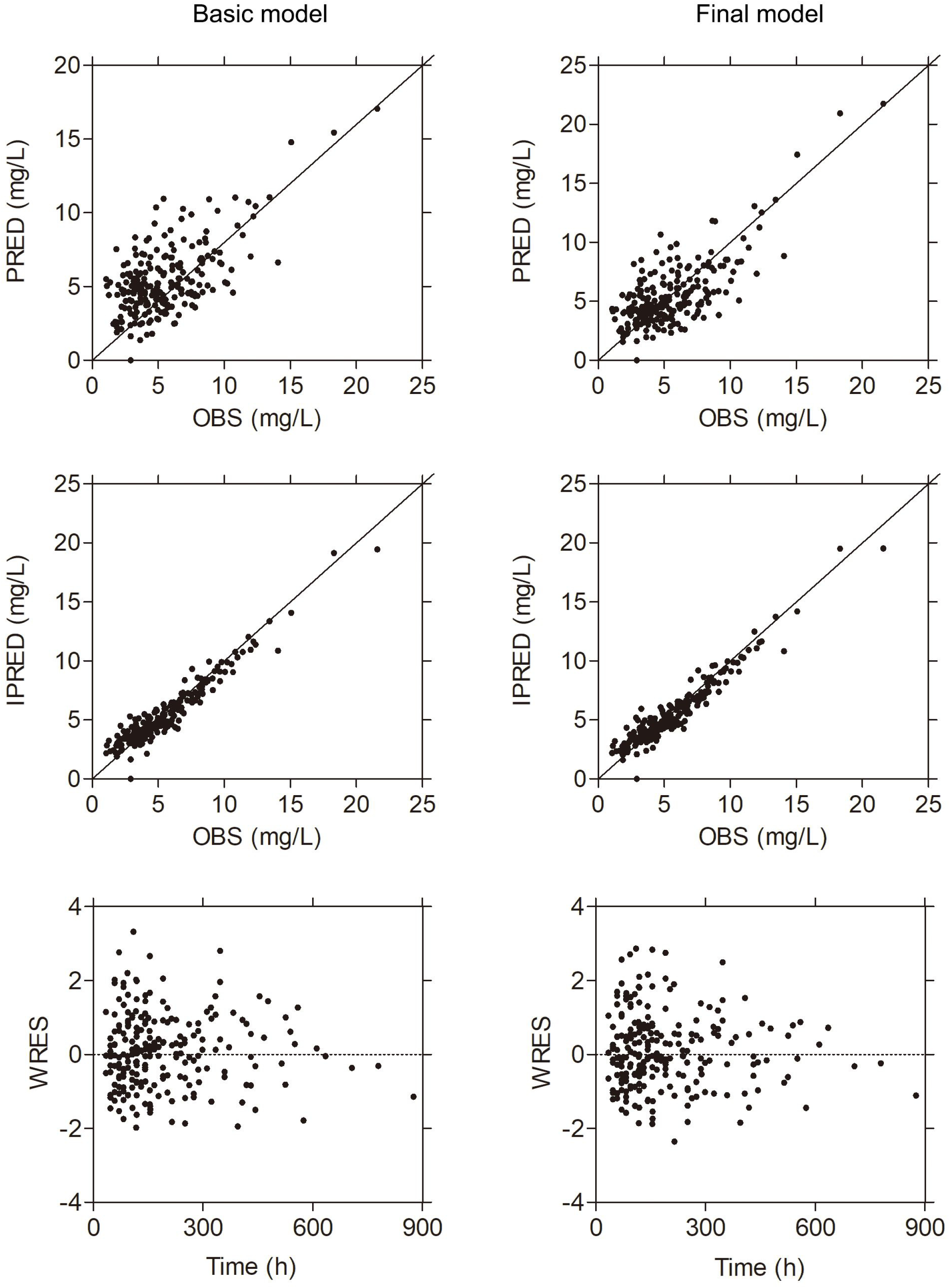
The goodness-of-fit of the basic model (left) and final model (right). OBS: observed concentrations; PRED, population predicted concentrations; IPRED, individual predicted concentrations; WRES: weighted residuals.

### 2.4 Simulations

The lack of effect of weight on voriconazole pharmacokinetics parameters does not support weight-adjusted dosing in patients with liver cirrhosis. Figure 3 and Figure S2 show the simulated concentration–time profiles between 0 h and 420 h calculated for the different dosage regimens. The probabilities of simulated voriconazole *C*_min_ values of <1 mg/L, 1–5 mg/L, and >5 mg/L with different oral and intravenous regimens in liver cirrhotic patients are presented in Table 4 and Table S1. The results of the simulations showed that the halved dose in CP-A/CP-B and CP-C patients could lead to a majority of cases for *C*_min_ over the supratherapeutic target (5 mg/L). Regarding the *C*_min_ target for efficacy and safety, the results of the simulations suggested that maintenance dosage regimens of 75 mg/12 h and 150 mg/24 h intravenously or orally for CP-A/CP-B patients, and of 50 mg/12 h and 100 mg/24 h intravenously or orally for CP-C patients are appropriate, since the predicted probability of achieving the therapeutic target concentration in the steady state was 59.0% – 68.2% for these optimal regimens in patients with liver cirrhosis.

**Table 4.** The probability of simulated voriconazole trough concentrations achieving or not achieving the therapeutic target for different oral dosage regimens in patients with liver cirrhosis.

| Dosage regimes | Probability (%) |  |  |  |  |  |  |  |  |
| --- | --- | --- | --- | --- | --- | --- | --- | --- | --- |
|  | Day 3 |  |  | Day 7 |  |  | Day 14 |  |  |
|  | < 1 | 1 - 5 | > 5 | < 1 | 1 - 5 | > 5 | < 1 | 1 - 5 | > 5 |
|  | mg/L | mg/L | mg/L | mg/L | mg/L | mg/L | mg/L | mg/L | mg/L |
| 100 mg/12 h, p.o.<br>for CP-A/ CP-B | 10.8 | 65.8 | 23.4 | 10.8 | 61.9 | 27.3 | 9.8 | 57.8 | 32.5 |
| 75 mg/12 h, p.o.<br>for CP-A/ CP-B | 13.7 | 68.2 | 18.1 | 15.0 | 66.8 | 18.2 | 15.2 | 64.9 | 19.9 |
| 150 mg/24 h, p.o.<br>for CP-A/ CP-B | 14.8 | 67.6 | 17.6 | 17.3 | 66.9 | 15.8 | 18.7 | 63.6 | 17.7 |
| 100 mg/12 h, p.o.<br>for CP-C | 5.8 | 53.0 | 41.3 | 4.2 | 45.4 | 50.4 | 2.8 | 36.5 | 60.7 |
| 50 mg/12 h, p.o.<br>for CP-C | 9.3 | 62.9 | 27.7 | 10.3 | 64.8 | 24.9 | 10.2 | 64.3 | 25.5 |
| 100 mg/24 h, p.o.<br>for CP-C | 9.4 | 63.5 | 27.1 | 11.3 | 64.5 | 24.2 | 11.6 | 64.0 | 24.4 |

CP-A/CP-B: Child-Pugh Class A and Child-Pugh Class B; CP-C: Child-Pugh Class C; p.o.: orally.

**Figure 3.**
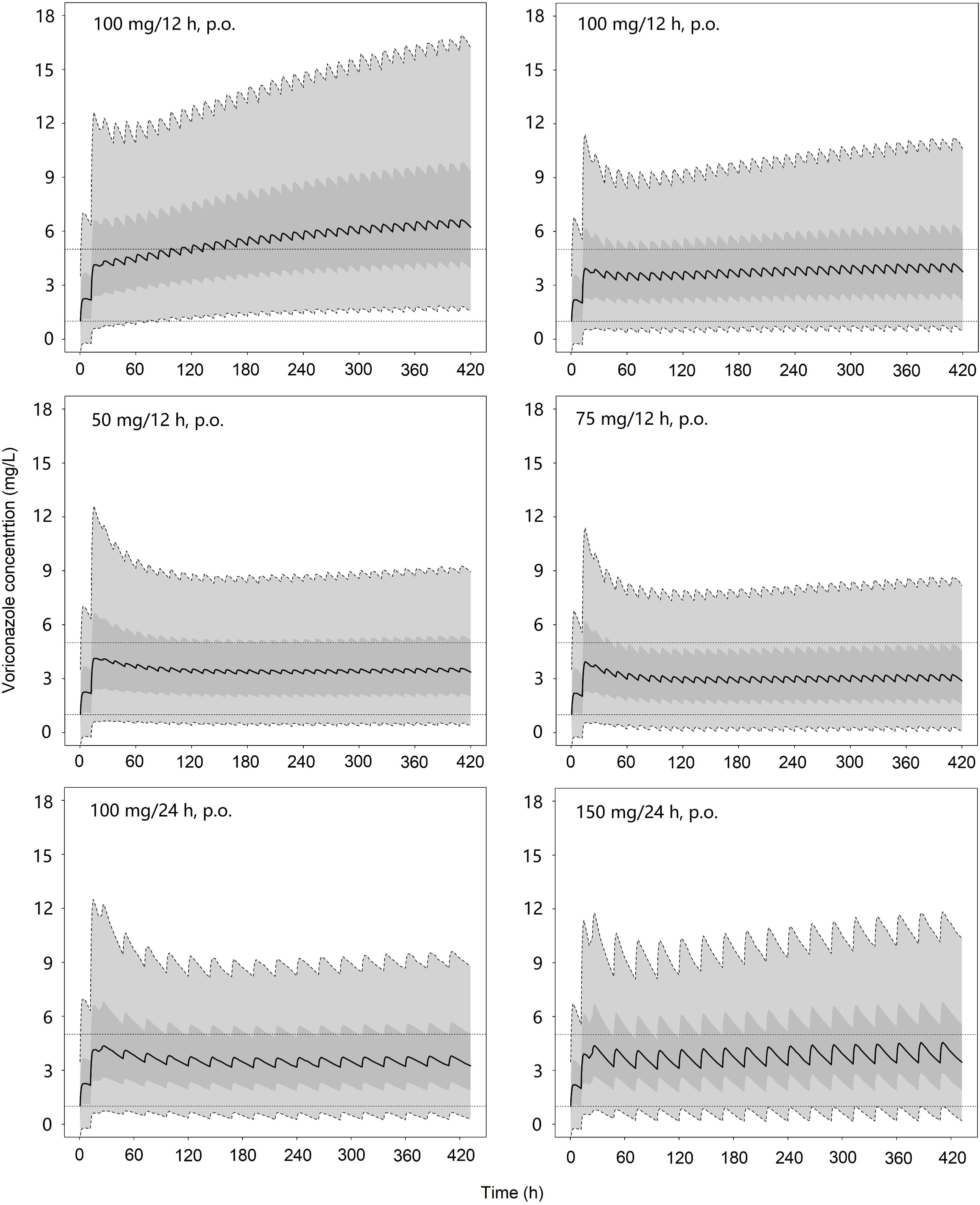
The concentration–time profiles of different voriconazole oral dosing regimens in CP-C (left) and CP-A/CP-B (right) patients based on 50000 simulations. The dashed lines represent the voriconazole target trough plasma concentrations of 1 mg/L and 5 mg/L. The solid line is the 50% of simulated data, and the dark grey area and the light gray area are the 50% (Q_25-75_) and 90% (Q_5-95_) prediction intervals, respectively.

## 3. Discussion

The rationality of halving the voriconazole dose in CP-A/CP-B patients has not been validated widely in clinical practice, and the optimal dose in CP-C patients remains to be established. The present study used a PPK analysis to identify the potential source of pharmacokinetics variability and to optimize the dosage regimens in patients with liver cirrhosis.

To our knowledge, this is the first time that voriconazole PPK properties have been characterized in patients with liver cirrhosis. In the present study, single- or multiple-trough samples were collected from patients. These features restricted the ability to estimate *k*_a_ and V_d_ accurately, which may have an impact on the simulated peak concentration of voriconazole. In the process of constructing the basic model, the OFV value of the model was 532.6 and 535.5, respectively, when *k*_a_ was fixed to a value of 0.5 h^−1^ (16) or 1.1 h^−1^ (17), as reported elsewhere. However, we found that the variabilities of *k*_a_ and *F* were very small when the NONMEM software was used to calculated *k*_a_ instead of giving a fixed value, and a OFV value of 508.6 was observed in the process. Thus, the basic model in this study did not include *k*_a_ and *F* variability, and a *k*_a_ value of 0.25 h^−1^ was calculated, which is similar to previously reported values (0.53 h^−1^ to 1.1 h^−1^) (16-18). Actually, delayed gastric emptying in cirrhotic patients could delay the absorption of drugs by the small intestine (19). The estimated bioavailability of voriconazole in this study was 91.8%, which was slightly higher than previously reported values for adult patients (63.0% – 89.5%) (17, 18, 20, 21). Theoretically, the reduced liver blood flow and enzyme activity in cirrhotic patients can substantially reduce the first-pass effect, which results in greater systemic availability (22).

Like the Child-Pugh class, the nine potential covariates (TBIL, Child-Pugh score, INR, MELD score, GGT, AST, ALP, PLT, and ALT) are also indicators of liver function, and so these nine covariates and the Child-Pugh class were not incorporated in a model simultaneously. Additionally, the simultaneous addition of Child-Pugh class and age did not lead to a further significant decrease in OFV. Thus, the Child-Pugh class was found to influence voriconazole CL and was the only covariate included in the final model. As the samples were trough values, the CL can be accurately estimated using PPK method. The estimated typical values of CL for CP-A/CP-B and CP-C patients were 1.79 L/h and 0.99 L/h, respectively, which suggest that different dosing regimens need to be administered depending on the Child-Pugh class. The calculated CL in cirrhotic patients is much lower than that in patients with normal liver function (17, 21). Actually, the Child-Pugh class is the most-common method used to classify liver impairment and to adjust the dose of drugs in patients with liver cirrhosis (23, 24), which mainly due to liver disease being clearly associated with reduced liver blood flow and enzyme activity (22). The elimination *t*_1/2_ value of voriconazole is significantly prolonged in patients with liver cirrhosis, and so we believe that it is sufficient to administer voriconazole once daily in cirrhotic patients. The final PPK model appeared to describe the data very well and to be reliably validated, making it adequate for further dosing simulations. The results of the simulations showed that 32.5 – 36.7% and 60.7 – 66.2% of CP-A/CP-B and CP-C patients, respectively, had a *C*_min_ exceeding the supratherapeutic target on at day 14 after they had received the halved dose (Table 4 and Table S1). Thus, the halved dose may be inappropriate in cirrhotic patients for an amount of supratherapeutic voriconazole *C*_min_ emerged. For CP-C patients, only Yamada *et al.* (25). has provided an optimal voriconazole maintenance dose in CP-C patients; those authors used clinical data from six CP-C patients and concluded that the oral voriconazole maintenance doses should be reduced to one-third of the normal dose. However, the results obtained in the present model-based simulations indicate that voriconazole maintenance doses in cirrhotic patients should be reduced to one-fourth for CP-C patients and to one-third for CP-A/CP-B patients compared to that for patients with normal liver function. These optimal dosage regimens could ensure that >60% of patients will achieve a *C*_min_ within the therapeutic target. Notably, TDM was also necessary in patients who received the optimal dosage regimens, since 20 – 30% of them exhibited *C*_min_ values that exceeded the supratherapeutic target. Furthermore, further clinical studies are needed to investigate the safety and efficacy of these optimal dosage regimens in adequate numbers of cirrhotic patients.

This study found a clear association between voriconazole *C*_min_ and AEs, and that the threshold *C*_min_ for identifying patients with a higher risk of AEs was 5.12 mg/L, which is similar to the supratherapeutic target of voriconazole treatment recommended by the guideline from the British Society for Medical Mycology (12). The rate of AEs was higher in this study than previously reported in liver transplant patients (26). A particularly interestingly finding was that 69.0% (20/29) of cases of voriconazole-related AEs developed within the first week after voriconazole treatment, and that the elimination *t*_1/2_ was prolonged, which suggested that implementing accurate initial dose or plasma concentration monitoring as early as possible could avoid the occurrence of AEs.

This study was subject to some limitations. First, it couldn’t evaluate the hepatotoxicity according to the criteria of the National Cancer Institute in cirrhotic patients. Additionally, this study had a retrospective design. These two factors would reduce the incidence of voriconazole-related AEs. Second, most of patients did not detect *CYP2C19* polymorphisms in this study. Our previous study showed that voriconazole had a *C*_min_ of 3.13 ± 2.68 mg/L in poor metabolizers of *CYP2C19* with normal liver function who received the recommended dosage regimens (27). This suggests that the effect of liver cirrhosis on voriconazole pharmacokinetics parameters may exceed the effect of the *CYP2C19* polymorphism. Third, the number of Child-Pugh Class A patient was too small (*n* = 6), and further studies are needed to optimize voriconazole dosage regimens in these patients.

## 4. Conclusion

We have firstly explored the PPK parameters of voriconazole in patients with liver cirrhosis, with the findings suggesting that the optimal voriconazole dosages are 75 mg/12 h and 150 mg/24 h intravenously or orally for CP-A/CP-B patients, and 50 mg/12 h and 100 mg/24 h intravenously or orally for CP-C patients. To reduce the incidence of voriconazole-related AEs，it is critical to ensure voriconazole *C*_min_ is <5.12 mg/L by monitoring voriconazole *C*_min_ early. Clinically, TDM is also needed in liver patients who receive the optimal dosage regimens because *C*_min_ exceeded the supratherapeutic target in some patients. The present study represents a substantial step forward in determining the optimal dosing regimens in cirrhotic patients, and should be complemented by future clinical studies.

## 5. Materials and methods

### 5.1 Patients

We reviewed the laboratory information system from May 2014 to May 2018 at two tertiary hospitals. Patients who treated by voriconazole and therapeutic drug monitoring (TDM) had been performed during antifungal therapy were identified and then screened by applying inclusion criteria of (1) presence of liver cirrhosis, including CP-A, CP-B and CP-C patients and (2) age >18 years, and exclusion criteria of (1) the initial TDM after dose adjustment and (2) missing the values of key covariates. This study was approved by all participating institutional review boards. Finally, we retrospectively collected 120 cirrhotic patients into the analysis. Among these patients, four initial dosage regimens were administered: (1) patients received the recommended dosage regimen based on package insert or a fixed dose of 200 mg twice daily orally or intravenously; (2) patients received the loading dose of 200 mg twice daily on day 1, following by 100 mg twice daily orally or intravenously or a fixed dose of 100 mg twice daily orally or intravenously; (3) patients received loading dose of 200 mg twice daily on day 1, following by 200 mg daily orally or intravenously or a fixed dose of 200 mg daily orally or intravenously; (4) patients received a fixed dose of 100 mg daily orally or intravenously.

### 5.2 Data collection

Data were extracted from laboratory information system and medical records, including voriconazole-related information (*C*_min_, timing, amount, route), demographics (age, gender and bodyweight), the type of cirrhosis, laboratory data (the liver, kidney and coagulation function) and concomitant medications (CYP2C19 inhibitors and inducers). The Child-Pugh score, Child-Pugh class and MELD (Model for end-stage liver disease) score were calculated based on previous criteria (28, 29). The types of AEs were extracted from medical records, and the causal relationship of AEs with voriconazole treatment was defined according to the CTCAE criteria over the treatment period (30).

### 5.3 Blood sampling and analytical assays

A *C*_min_ was defined as a level obtained after 3 days of voriconazole therapy with a loading dose or 5 days of voriconazole treatment without a loading dose, and samples were collected at interval windows of 10 – 12 h post-dose. The validated high-performance liquid chromatography (HPLC) assays from participating hospital were used to measure voriconazole concentrations (9, 31). We use blank plasma to dilute the plasma whose concentration was beyond the upper limit of quantification and then measured by HPLC.

### 5.4 Population pharmacokinetics model and simulations

#### 5.4.1 Population pharmacokinetics model construction

A population pharmacokinetics (PPK) analysis was conducted using the data to estimate the typical values and the variabilities of pharmacokinetics parameters in the population, and to identify the factors influencing these parameters.

Firstly, a basic model, including structural model and statistical model, was developed to describe the PK properties of voriconazole without consideration of covariate effects. A one compartment model with first-order elimination was found to be appropriate as the structural model. The pharmacokinetics parameters, clearance (CL), volume of distribution (V_d_), absorption rate constant (*k*_a_) and bioavailability (*F*) were calculated in this model. An exponential model described the inter-individual variability with a mean of zero and a variance of ω^2^, and a combined error model described residual variability with a mean of zero and a variance of σ^2^. The elimination half-life, *t*_1/2_, was calculated by the form of ln(2)/(CL/V_d_).

Next, the effect of each covariate on a single pharmacokinetics parameter was tested in a single functional form separately, with a reduction of the objective function value (OFV) greater than 3.84 considered statistically significant. These retained covariates were then used to construct the PPK model in the forward selection process. A backward elimination process was not used in this study since only one covariate was incorporated in the forward selection process. All of these processes were followed the standard procedure of model building (32). The most appropriate PPK model had to meet the following criteria: (1) the minimal OFV; (2) A reduction in OFV of > 6.83 in the process of forward selection; (3) the diagnostic plots were improved compared to the basic model. (4) the covariate should be physiologically plausible.

#### 5.4.2 Model validation

The precision of voriconazole parameters and the robustness of the final model were evaluated using non-parametric bootstrap. A total of 1000 runs were carried out. The visual predictive check (VPC) was used to evaluated the predictive performance of the final voriconazole PPK model. A total of 1000 replicates were simulated, and the 90% prediction intervals were generated. The simulated results were compared with these observed concentrations in the present study visually.

#### 5.4.3 Model-based simulations for optimizing voriconazole doses

The final PPK model was used to simulate the voriconazole plasma concentration–time profile over 420 h by considering the variability of pharmacokinetics parameters and covariate. This was used to generate 50,000 pharmacokinetics profiles of voriconazole for each candidate regimen in CP-A/CP-B and CP-C patients, and the dosing schedules were set to 30, 50, 75, or 100 mg twice daily orally or intravenously, and 60, 100, 150, or 200 mg daily orally or intravenously. All of the simulated virtual patients received a loading dose of 400 mg twice daily orally or intravenously. The concentration–time profiles of the simulated virtual patients were summarized and compared using their median and 5%, 25%, 75%, and 95% percentile values. These simulations aimed at predicting the achievement of the voriconazole therapeutic target of 1 – 5 mg/L (12).

In the present study, PPK analysis and model-based simulations were performed using NONMEM software (version 7.20) and subroutine ADVAN6 and the first-order conditional estimation with interaction algorithm, and Wings for NONMEM (Version 743). The voriconazole plasma concentration-time curve were drew with R software (version 3.4.4).

### 5.5 Statistical analysis

The Child-Pugh class was included as a dichotomous variable in this study, with a value of 0 indicating CP-A/CP-B patients and a value of 1 indicating CP-C patients. Student’s *t*-test was used to compare voriconazole troughs. A classification and regression tree (CART) analysis was used to identify the threshold for a continuous variable that would maximize the difference in outcome between two groups. A *P* value of < 0.05 was considered statistically significant.

## Transparency declarations

None to declare.

## Ethical approval

This study was approved by all participating Hospital Ethics Committee.

